# Cryo-electron tomography reveals postsynaptic nanoblocks in excitatory synapses

**DOI:** 10.1101/2023.05.12.540562

**Authors:** Rong Sun, James P Allen, Liana Wilson, Mariam Haider, Baris Alten, Zimeng Zhou, Xinyi Wang, Qiangjun Zhou

**Affiliations:** Department of Cell and Developmental Biology, Center for Structural Biology, Vanderbilt Kennedy Center, Vanderbilt University, Nashville, TN, 37240, USA; Vanderbilt Brain Institute, Vanderbilt University, Nashville, TN, 37240, USA; Center for Structural Biology Cryo-EM Facility, Vanderbilt University, Nashville, TN, 37240, USA; Department of Pharmacology, Vanderbilt University, Nashville, TN, 37212, USA; School of Engineering, Vanderbilt University, Nashville, TN, 37212, USA; Peabody College, Vanderbilt University, Nashville, TN, 37212, USA; Harvard Medical School, Boston, MA, 02115, USA and Department of Neurology, Massachusetts General Hospital and Brigham & Women’s Hospital, Boston, MA, 02115, USA

## Abstract

The nanoscale organization of proteins within synapses is critical for maintaining and regulating synaptic transmission and plasticity. Here, we use cryogenic electron tomography to directly visualize the three-dimensional architecture and supramolecular organization of pre-cleft-postsynaptic components in their near-native cellular context in both synaptosomes from rat hippocampi and synapses from rat primary cultured neurons. High-resolution electron microscopy and quantitative analyses revealed that postsynaptic density (PSD) is composed of membrane-associated nanoblocks of various sizes, which are in close relationship with potential presynaptic release sites through adhesion molecules spanning the synaptic cleft, as well as with post-synaptic receptors. Subtomogram averaging from synaptosomes showed two distinct types of postsynaptic membrane proteins at resolutions of 24 Å and 26 Å respectively. Furthermore, our data reveal that majority of potential release sites and ∼50% subtomogram averaged receptor-like particles are located within the boundary of PSD nanoblocks, while PSD nanoblocks might be redundant for neurotransmission. The results of this study provide a more comprehensive understanding of synaptic ultrastructure and suggest that PSD is composed of clustering of various nanoblocks, which likely underlies the dynamic nature of PSD to modulate synaptic strength.

## Introduction

Cellular function is driven by nanoscale spatial organization of multi-protein assemblies and their interactions with other molecular components and cellular organelles. Typical examples of such nanoscale structures are neuronal chemical synapses. These are the most complex cell-cell junctions, with more than 2,000 distinct synaptic proteins^1^. Compelling evidence suggests that that multiple presynaptic, postsynaptic, and cell-adhesion proteins in excitatory synapses are finely organized in lipid-bound nanoscale structures to facilitate extremely rapid, dynamic, efficient, and tightly regulated transmission of information between neurons^2–10^. Moreover, the activity-dependent rearrangement of these synaptic proteins, including neurotransmitter receptors and other synaptic proteins, plays a key mechanistic role in synaptic plasticity, which is essential for perception, decision making, learning, and memory formation.

Recent studies provide evidence for a high-level organization of nanoscale supramolecular assemblies spanning the synaptic cleft, giving rise to *nanocolumns* of juxtaposed pre- and postsynaptic compartments in excitatory synapses^3,4,11–16^. Specifically, presynaptic active-zone scaffold proteins, such as RIM, form nanodomains that define synaptic vesicle docking and release sites. These sites align with clustered neurotransmitter receptors such as α-amino-3-hydroxy-5-methyl-4-isoxazolepropionic acid receptor (AMPAR) and postsynaptic scaffolding proteins such as PSD95 via adhesion proteins in the synaptic cleft^10,17,18^.

The postsynaptic density (PSD) appears in conventional electron micrographs as an electron-dense lamina just beneath the postsynaptic membrane^19^. A PSD meshwork model comprising regularly spaced vertical filaments was proposed based on electron microscopy (EM) and biochemical assays^20–24^. Two additional forms of PSD organization have been proposed: nano-domains based on super-resolution imaging^3–5,25–27^, and liquid condensate based on *in vitro* PSD mixing assay^28–30^. All these studies provide more evidence for the high-level organization of nanoscale supramolecular assemblies in the synapse, indicating that synaptic proteins are not randomly distributed and instead form functional synaptic nanostructures. However, nanoscale organization of the PSD and postsynaptic receptors is largely unknown due to the technical challenges imposed by synaptic molecular complexity.

Cryo-electron tomography (cryo-ET) is a powerful technique that produces three-dimensional (3D) volume reconstructions, or *tomograms*, of macromolecular and subcellular biological structures in their native context at nanometer resolution. To prevent introducing artifacts through chemical fixation, dehydration, and heavy-metal staining, biological samples are rapidly frozen to obtain a “true, instantaneous snapshot”^31–34^. In general, obtaining high-quality data requires cryo-ET samples to be thin enough for electrons to pass through (∼200 nm), posing a technical challenge during sample preparation, which limits the resolution of cellular cryo-ET^35^. The resolution is also limited by the crowded cellular environment. To overcome these limitations, isolated synaptic terminals have been used for cryo-ET studies to decrease sample thickness^10,36^.

A classic type of isolated synaptic terminal called a *synaptosome* is highly suitable for cryo-ET, because it retains a piece of attached postsynaptic membrane that faces the active zone^37,38^. Previous studies have demonstrated that PSD complexes and PSD membranes are resistant to detergents and stable enough to be purified from conventional synaptic fractions fractions^39–42^. This stability suggests that the core scaffold of the native PSD is likely maintained on the attached postsynaptic membrane^42^. Consequently, synaptosomes are ideal for cryo-ET data collection at high resolution, as the average thickness of a synaptosome is 200-300 nm in our preparations. Here, we developed a simplified sample preparation workflow to obtain the synaptosomes from mature rat hippocampi, achieving high-resolution synaptic nanostructures with cryo-ET and sub-tomogram averaging. With this method, we directly visualized the overall architecture of the transcellular organization in excitatory synapses. Our results revealed that PSD is formed by subsynaptic PSD nanoblocks of specific sizes. We also determined structures of two types of postsynaptic membrane proteins at 24-Å and 26-Å resolution.

## Results

### Transsynaptic organization of synaptosomes

To study subsynaptic organization, we employed a simplified workflow to prepare synaptosomes. After dissecting the hippocampi from rat brains, we performed four strokes of homogenization and a 10-minute centrifugation at 800 *g,* followed by plunge-freezing and cryo-ET data acquisition (Supplementary Fig. 1a). Our preparations yielded two different types of isolated synaptic terminals based on their morphology: synaptosomes retained a patch of postsynaptic membrane with PSD (Fig. 1 and Supplementary Fig. 1b-e, see also Supplementary Video 2), while synaptoneurosomes retained an enclosed postsynaptic compartment (Supplementary Figs. 1f-i and 2g)^43,44^. For this study, we primarily focused on synaptosomes, since they can provide higher-resolution structural information compared to synaptoneurosomes and intact synapses due to lower samples thickness less molecular crowding. In total, we acquired 123 tomograms of isolated synaptic terminals from three preparations without protease inhibitor or DTT in the homogenization buffer (99 synaptosomes and 24 synaptoneurosomes). This allowed us to directly visualize characteristic features of a synapse, including typical synaptic proteins (Fig. 1a, 1b and Supplementary Fig. 1p-u). In all our preparations, we successfully identified both synaptosomes and synaptoneurosomes, and no noticeable morphological differences were observed at the nanoscale across different batches of sample preparations. All the synaptosomes we obtained were likely from excitatory synapses, as they have thick PSDs of more than 30 nm, which is a known feature of excitatory synapses^19,22^.

**Fig. 1.**
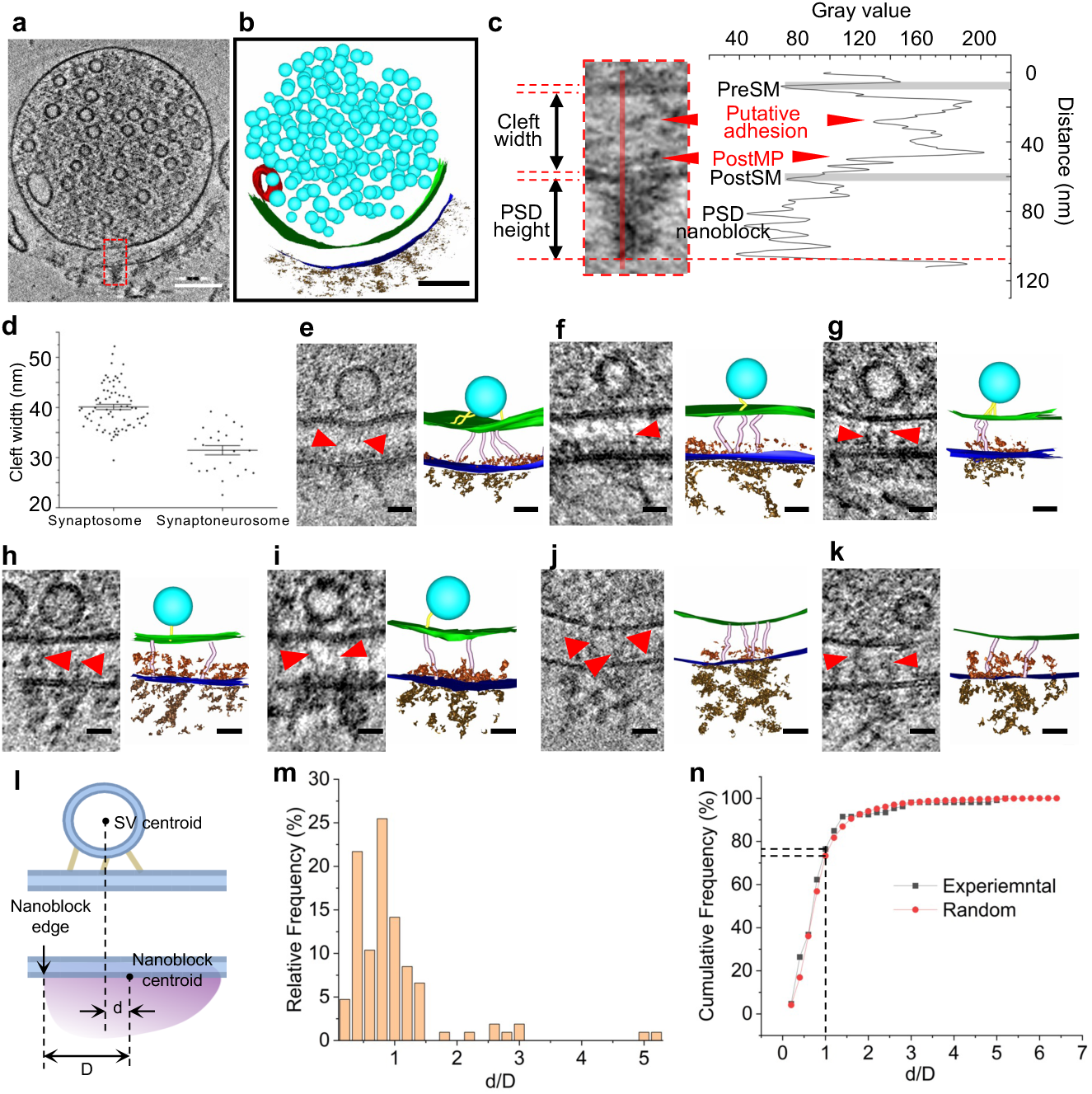
A potential presynaptic release site links with adhesion molecules, membrane proteins, and PSD nanoblocks. (a) A tomographic slice showing an excitatory synaptosome. (b) 3D segmentation of the synaptosome shown in panel a. (c) Quantification of the synaptic cleft and the PSD shown in panel a. The zoomed-in tomographic slice showing the excitatory synaptosome from panel a is on the left and its density plot showing gray values along the red solid line is on the right. (d) Cleft widths of synaptosomes and synaptoneurosomes. (e-k) Examples of transsynaptic alignment of adhesion molecules and postsynaptic membrane proteins in the cleft, as well as PSD nanoblocks in the postsynapse. Tethered synaptic vesicles also align with cleft and PSD densities. Red arrowheads indicate putative adhesion molecules in the cleft. (l) A schematic showing distance between synaptic vesicle centroid and nanoblock centroid, and distance between nanoblock centroid and edge. (m-n) Distribution of normalized distance between potential release site and nanoblock. d/D≤1 indicates that the SV centroid aligns within the PSD nanoblock. Scale bars: (a-b) 100 nm; (e-g) 20 nm. *Abbreviations PreSM: Presynaptic membrane. PostSM: Postsynaptic membrane. PostMP: Postsynaptic membrane protein*.

To study the relationship between potential release sites, adhesion molecules, postsynaptic densities, and postsynaptic receptors, we examined the tethered, docked or fused synaptic vesicles that likely indicate the location of synaptic release sites. We also investigated the corresponding nanostructures in the cleft and in the postsynaptic compartment. Examining the electron densities and measuring the gray values of synaptosome tomograms (Fig. 1c) allowed us to detect the adhesion molecule-like proteins and membrane proteins in the synaptic cleft, as well as the PSD protein clusters, which we refer as *nanoblocks*. The tomographic slice in Fig. 1c’s left panel shows protein densities in the middle of the cleft, which we posit to be the adhesion molecules, as well as a density layer on the postsynaptic membrane that is comprised of membrane proteins. These observations based on direct visualization are presented with the density plot in Fig. 1c’s right panel. We found that synaptosomes and synaptoneurosomes have specific cleft widths (Fig. 1d); and the clefts of synaptosomes tend to be wider than those of synaptoneurosomes (40.1±0.5 nm for synaptosomes n=74; and 31.5±0.9 nm for synaptoneurosomes n=24). Of note, 15 synaptosomes were not included in these measurements as the postsynaptic membranes were not parallel to the presynaptic membranes. The disparity in cleft widths between synaptosomes and synaptoneurosomes may result from distinct adhesion molecules that withstand the sample isolation process. We observed that tethered vesicles are in close proximity of the adhesion-like molecules, the membrane protein-like particles, and PSD densities (Fig. 1e-i). This direct visualization of pre-cleft-postsynaptic structure suggests that PSD scaffold proteins, might be specifically aligned with potential presynaptic release sites through adhesion molecules in the synaptic cleft. Two examples of adhesion-PSD nanoblock alignment without a tethered synaptic vesicle are also shown here, indicating these might be potential unoccupied release sites (Fig. 1j-k). To quantify the relationship between PSD nanoblock and potential release sites occupied by docked, tethered, or fused synaptic vesicles, we analysed the distance from vesicle centroid to nanoblock centroids, normalized by the distance between nanoblock centroid to nanoblock edge, and compared with what is observed with random distribution. We found that 76.4% of potential release sites were transsynaptically located in the boundary of a nanoblock (Fig. 1l-n). However, considering all docked, tethered, or fused synaptic vesicles (n=106) in synaptosomes, the alignment between measured positions of synaptic vesicles and nanoblocks did not show significant difference from the alignment between randomized positions of synaptic vesicles and nanoblocks (Fig. 1n), which suggests that PSD nanoblocks are redundant for alignment with the potential release sites occupied by synaptic vesicles.

### PSD subsynaptic organization in synaptosomes

Next, we investigated PSD densities and performed quantitative analyses on the PSD nanoblock dimensions. A representative tomogram of a synaptosome acquired by cryo-ET is shown in Fig. 2a, in which PSD nanoblocks (indicated by red arrows) are clearly visualized beneath the membrane. In terms of the overall PSD EM density patterns, this result is consistent with previous EM studies stating that the core of the PSD is dominated by vertically oriented filaments^20,45^ or spiky structures^24^. Our data revealed three types of synaptosomes based on the organization of the PSD: synaptosomes with separated PSD nanoblocks; synaptosomes with both separated and continuous PSD nanoblocks; and synaptosomes with continuous PSD (Supplementary Fig. 1j-o). Importantly, it is evident that the distribution of PSD densities is not uniform across these three types of synaptosomes. The presence of PSD nanoblocks is clearly observable in the original 3D tomograms (Supplementary Fig. 1jo). To delineate the PSD nanostructures of all three synaptosome types, we applied the following quantitative analysis: First, we measured gray values of protein densities of a 40-nm thick PSD area beneath the postsynaptic membrane (Fig. 2a-b). Then, we combined 2D gray values across a series of continuous virtual tomographic slices, generating a gray value density map (Fig. 2c, top panel). With normalization and filtering processing, **<u>d</u>**ensity-**<u>b</u>**ased **<u>s</u>**patial **<u>c</u>**lustering of **<u>a</u>**pplications with **<u>n</u>**oise (DBSCAN) results show projected PSD density nanoblocks in the 40-nm range from the postsynaptic membrane (Fig. 2c, bottom panel). DBSCAN is a density-based clustering non-parametric algorithm^46^ and it is widely used in biological data analysis^47,48^. In this way, 2D DBSCAN results represented PSD densities flattened and projected to *xz-plane* (Fig. 2d). We next measured the area and width of each protein nanoblock (Fig. 2c and 2e). The area of nanoblock was defined as its convex hull area shown by encircling red lines. The width of nanoblock was defined as the largest *x-axis* distance between two pixels in each cluster. Fig. 2e shows examples of nanoblocks with different areas and widths quantified by the DBSCAN clustering method. Multiple peaks were detected in the area distribution of nanoblocks from all synaptosomes (Fig. 2f), while the width distribution was fit with two peaks (Fig. 2g). Distinct fitting peaks in the area statistics histogram indicate that nanoblocks have distinct sizes, and that large nanoblocks may be composed of several small nanoblocks, which is also supported by examining the morphology of some nanoblocks with large widths (Fig. 2e). Though the geometric shape of the nanoblock was irregular, the way we quantified the width reflected the dimension of the nanoblock in *x-axis*, while the dimension of nanoblock in *z-axis* was affected by the missing wedge effect^49^. We also tested DBSCAN with different parameters (Supplementary Fig. 2a-b) and DBSCAN (6, 32) was applied as the results were more consistent with the visualized electron density clusters in the tomographic slices. We isolated synaptosome sample from three separate preparations and obtained consistent cleft with and nanoblock sizes across all samples (Supplementary Fig. 2c-d). To further rule out the influence of sample preparation on these observed ultrastructural features, we collected an additional 83 tomograms of isolated synaptic terminals (72 synaptosomes and 11 synaptoneurosomes) from three preparations with protease inhibitor and DTT in the homogenization buffer. The data of synaptosomes treated with protease inhibitors and DTT (Supplementary Fig. 2e-f), along with the data of synaptoneurosomes (Supplementary Fig. 2g-h), collectively present compelling evidence that supports the presence of nanoblocks across different sample types, ruling out the possibility of isolation artifacts. Overall, our analyses suggest that PSD nanoblocks can be positioned closely enough to one another to form larger nanoblocks.

**Fig. 2.**
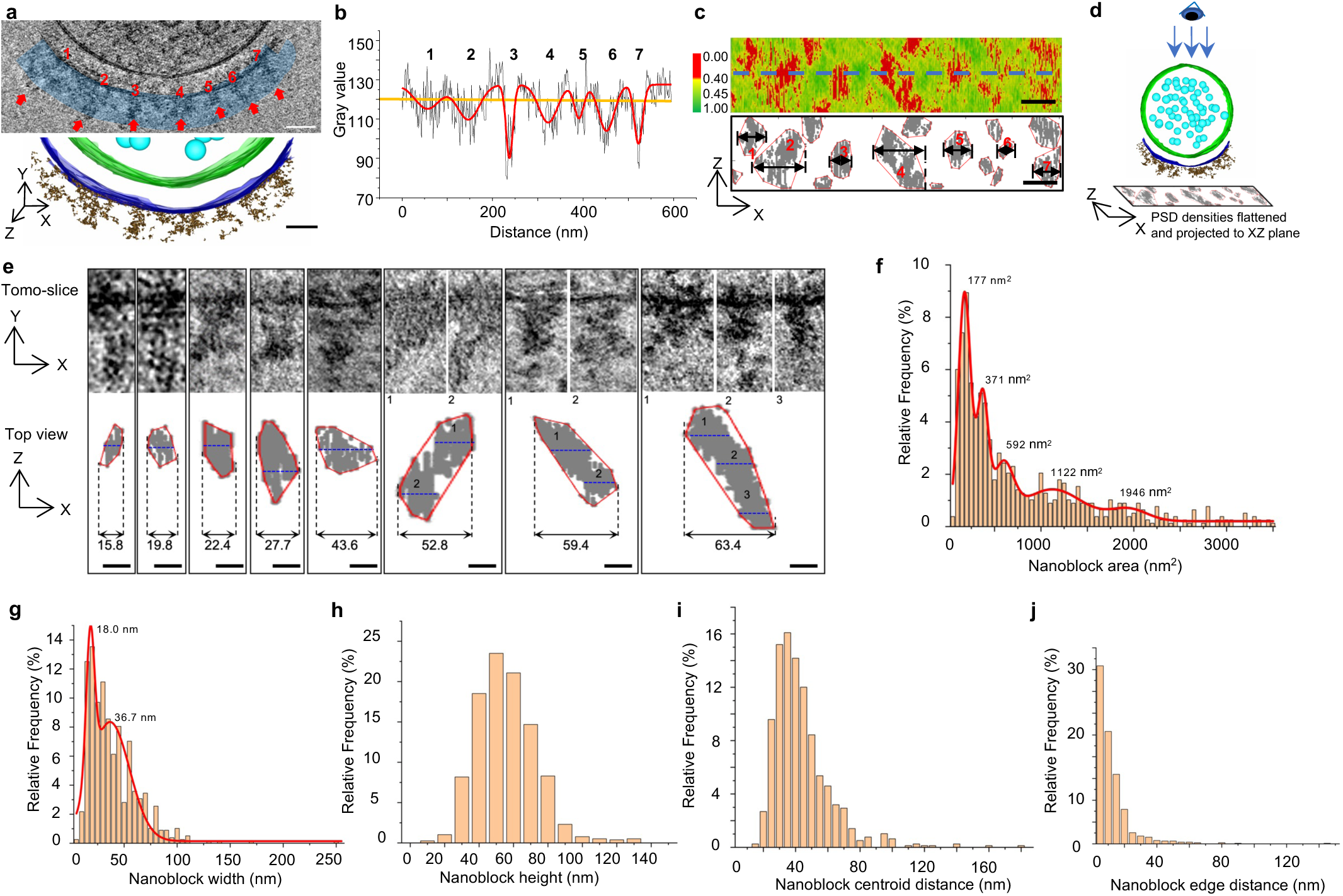
PSD subsynaptic organization in synaptosomes. (a) Top panel: A tomographic slice showing the subsynaptic nanoblocks of a synaptosome. Arrows and numbers indicate the seven nanoblocks in the PSD. Bottom panel: 3D segmentation of the synaptosome in the top panel. (b) Density plot shows average gray values (black lines) along the 40-nm blue bar in the top panel of (a). Red lines indicate the fitting curve of average gray values. (c) 3D density analysis in the PSD of synaptosome in (a). Top panel: Density plot shows gray values of 40-nm PSD density projected to the xz plane. The blue dotted line indicates the plane of the tomographic slice shown in (a). Bottom panel: DBSCAN clustering result of 40-nm PSD density projected to the xz plane. (d) A schematic showing the clustering results of PSD densities produced from 3D synaptosome data. (e) Examples of PSD nanoblocks in 2D tomographic slices and corresponding projected DBSCAN clustering results. Values in each panel indicate the width (Units: nm) of the PSD block. The blue dotted lines indicate the plane of the corresponding tomographic slices. (f) Distribution of PSD nanoblock area. (g) Distribution of PSD nanoblock width. (h) Distribution of nanoblock height. (i) Distribution of nearest neighbor distance between PSD nanoblock centroids. (j) Distribution of distance between nearest pixels of PSD nanoblocks. Scale bars: (a, c) 50 nm; (e) 20 nm.

To further analyse the size and morphology of subsynaptic nanoblocks, we measured the height of the protein nanoblocks obtained via DBSCAN clustering. The height was quantified based on gray values along the nanoblock at its largest length (Fig. 1c). The distribution of nanoblock heights revealed that most nanoblocks are in the 30∼90 nm range, while a small population is below 20 nm or above 100 nm (Fig. 2h). Additionally, we analysed the gap between PSD nanoblocks. The distance between each nanoblock centroid (Fig. 2i) and the nearest distance between each nanoblock edge (Fig. 2j) together describe the spacing between nanoblocks, suggesting that the gaps with less protein densities segmented the PSD. Overall, our analysis demonstrates that subsynaptic nanoblocks of different sizes form the PSD.

### Organization of membrane protein particles on the postsynaptic membrane

To further investigate the nanoscale organization of the PSD, we employed the subtomogram averaging method to detect postsynaptic membrane proteins in our synaptosome samples. We manually selected 1,565 protein particles from 28 synaptosomes in the range from 10 nm to 14 nm for subtomogram averaging (Fig. 3a and Supplementary Fig. 3a). Eventually, we achieved an averaged structure of type A particle with 391 particles (Fig. 3b) and an averaged structure of type B particle with 189 particles (Fig. 3c).

**Fig. 3.**
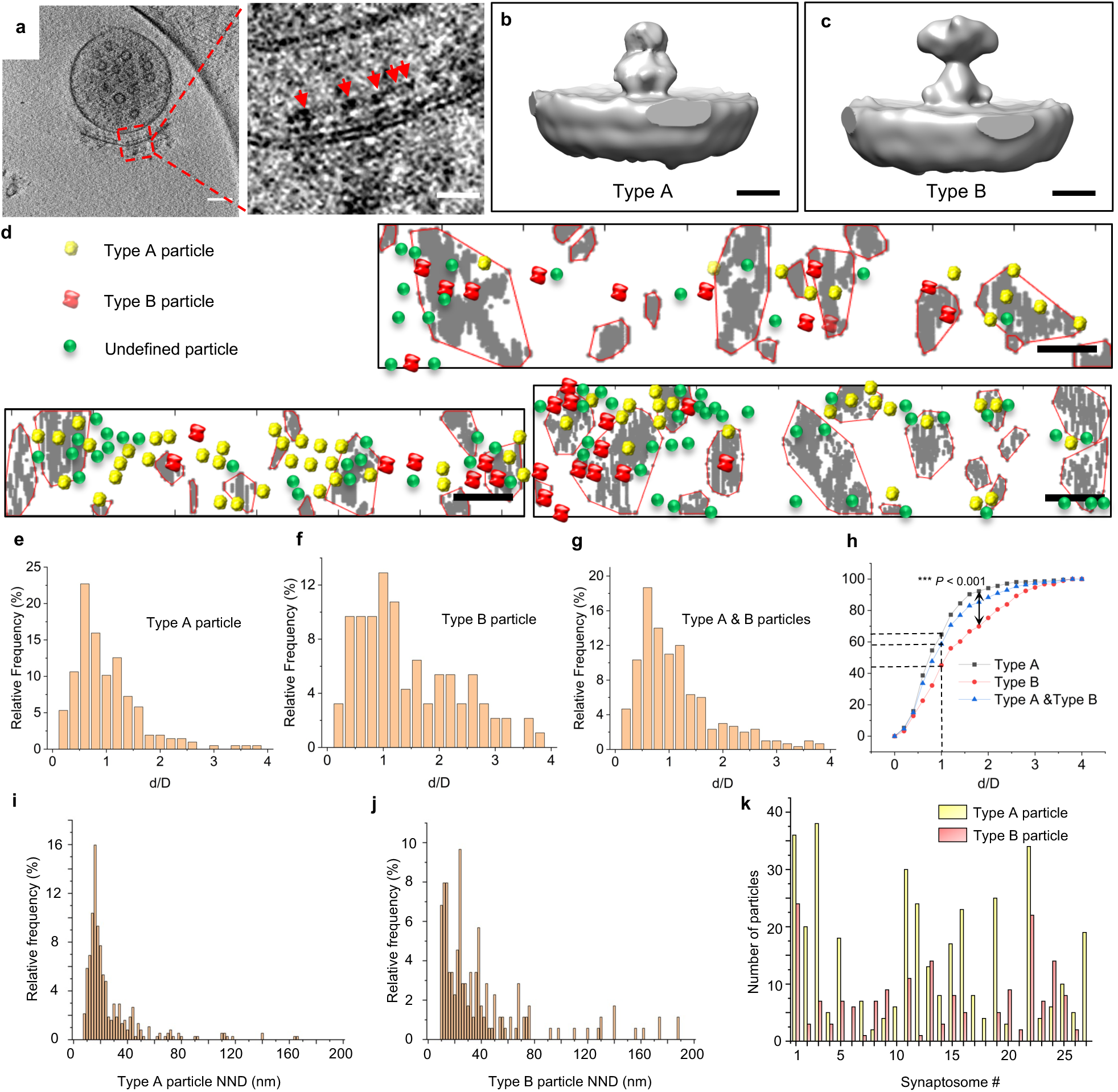
Subtomogram averaging of postsynaptic membrane particles. (a) Membrane particles on the postsynaptic membrane. (b-c) Subtomogram averaged type A and B particles. (d) Examples of type A, type B, and undefined particles on the postsynaptic PSD nanoblocks. (e-h) Distribution of normalized distance between type A or type B particles and nanoblock. (i-j) Nearest neighbor distances of type A particle (j) and type B particle (j). (k) Type A and type B particle numbers in each synaptosome. Scale bars: (a) left panel 100 nm, right panel 20 nm; (b-c) 5 nm; (d) 50 nm.

To study the postsynaptic distribution of the membrane particles, we superimposed the type A particles, type B particles, and all remaining undefined particles with the PSD clustering results (Fig. 3d). To quantify the relationship between these two types of membrane particles and nanoblocks, we used the same method with which we analysed potential release sites (Fig. 3e-f). This analysis was performed in 13 synaptosomes which were used both in PSD nanoblock quantification and subtomogram averaging. We found that 64.7% type A particles and 45.2% type B particles are within nanoblocks (d/D≤1). Additionally, we found distribution of d/D for type A and type B particles are different (two-sample Kolmogorov–Smirnov test, D=0.2551, n=207 for type A particles and n=93 for type B particles, p=3.54×10^-4^) and type A particles are closer to nanoblocks compared to type B particles (two-sample t test, n=207, mean=0.89 nm for type A particles and n=93, mean=1.34 nm for type B particles, p=2.52×10^-5^). We performed a nearest neighbor distance analysis and found type A particles are more clustered than type B particles (Fig. 3i-j, two-sample t test, mean for type A is 30.8 nm and mean for type B is 50.5 nm, p=0.00139). The number of type A and type B particles vary in all synaptosomes (Fig. 3k), showing the heterogeneity of the synaptosomes. However, the relatively low resolution achieved through subtomogram averaging renders it challenging to determine the protein identity of type A or type B particles with certainty.

### PSD subsynaptic organization in intact neurons

We also examined the subsynaptic organization of synapses to determine whether similar subsynaptic nanoblocks identified in synaptosomes can also be found in intact synapses from cultured neurons. To accomplish this, we seeded dissociated primary hippocampal neurons on gold EM grids and examined the synaptic ultrastructure in intact neurons using cryo-ET (Supplementary Fig. 4a-e). In reconstructed tomograms, a typical synapse consists of a presynaptic compartment with a population of synaptic vesicles, a synaptic cleft, and a postsynaptic compartment with PSD. We also focused on excitatory synapses and excluded synapses with thin PSDs in cultured neurons from our analysis (Supplementary Fig. 4f-h).

We collected 211 tilt series of synapse-like structures from the wild-type primary hippocampal neurons and reconstructed them. 131 of the reconstructed tomograms were synapses, and 121 synapses had thick PSDs. Three examples of excitatory synapses are shown in Fig. 4. Tomographic slices (Fig. 4a, c, f, h, k, and m) and segmented structures (Fig. 4b and g, see also Supplementary Video 2) display EM densities within the PSD area are not uniformly distributed, suggesting that protein distribution is not homogeneous within the PSD. However, the crowded molecular environment and overall higher thickness of intact neurons rendered a low signal-to-noise ratio, making patterns of protein complexes difficult to recognize and analyse. Using the software package IsoNet^50^, we performed contrast transfer function (CTF) deconvolution and corrected the missing wedge effect to enhance contrast in 34 out of 121 cryo-electron tomograms of excitatory synapses. Following IsoNet correction, tomographic slices show more robust protein nanoblocks in the PSDs (Fig. 4d, i, and n). Quantitative characterization of the PSDs’ 3D volume (Fig. 4e, j, and o) indicates that proteins distribute unevenly within the PSD and form subsynaptic nanoblocks. The size and distribution statistics of the PSD nanoblocks (Fig. 4p-r) closely resemble those obtained from synaptosomes and synaptoneurosomes (Fig. 2f and Supplementary Fig. 2e-h). Our observations suggest that intact synapses also feature nanoblocks of varying sizes composed of postsynaptic proteins, similar to those observed in synaptosomes and synaptoneurosomes.

**Fig. 4.**
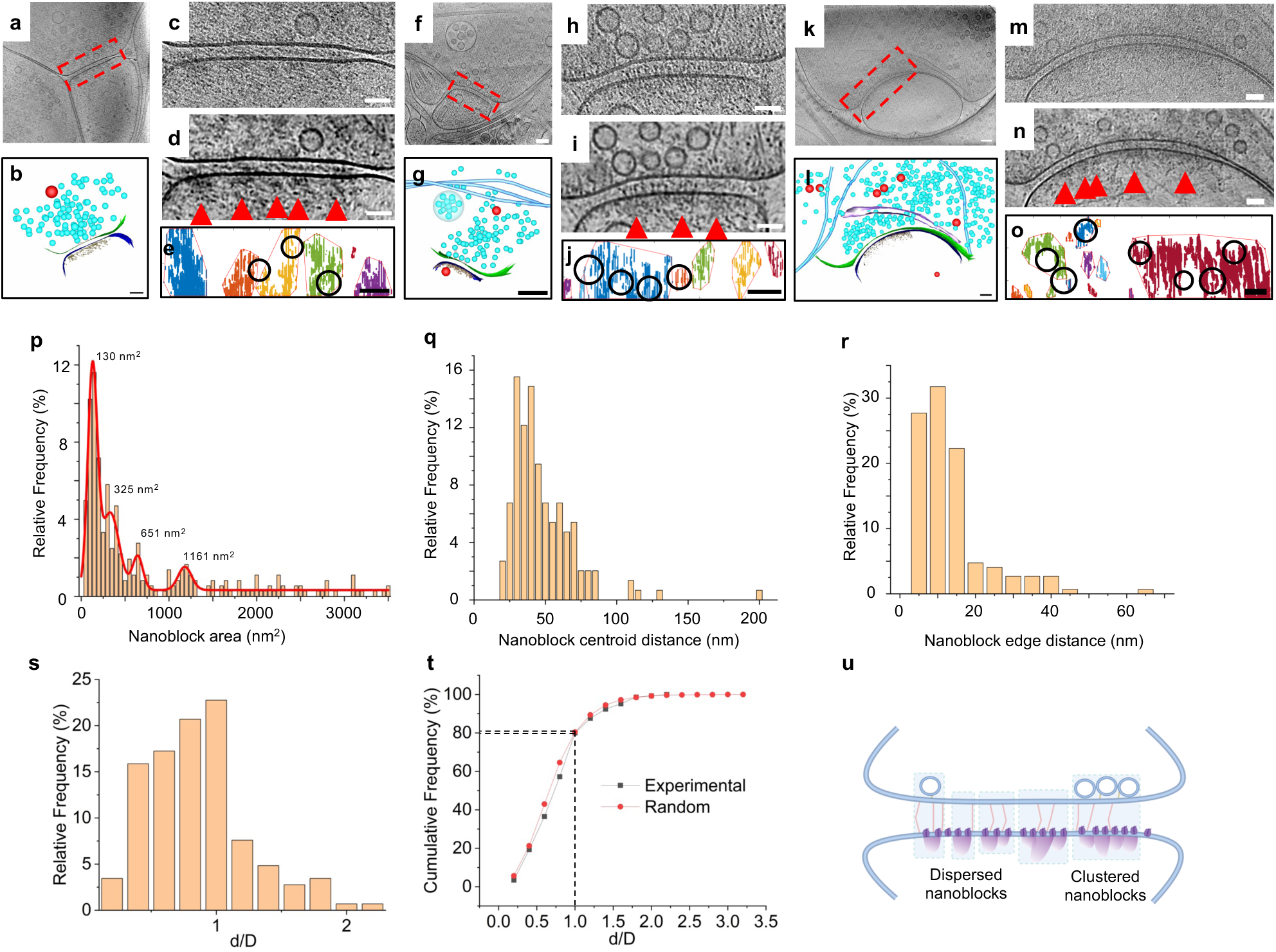
PSD subsynaptic organization in intact neurons. (a) Tomographic slice of an excitatory synapse. (b) Segmentation of the synapse in (a). (c) Zoom-in view of the synapse in (a). (d) IsoNet-corrected tomographic slice of (c). Red arrowheads indicate PSD nanoblocks. (e) Clustering results of 40-nm PSD density projected to the plane parallel to the postsynaptic membrane. Each color represents one cluster. Docked or tethered vesicle positions are marked as circles. (f) Tomographic slice of a second excitatory synapse. (g) Segmentation of the synapse in (f). (h) Zoom-in view of the synapse in (f). (i) IsoNet-corrected tomographic slice of (h). Red arrowheads indicate PSD nanoblocks. (j) Clustering results of 40-nm PSD density projected to the plane parallel to the postsynaptic membrane. Each color represents one cluster. Docked or tethered vesicle positions are marked as circles. (k) Tomographic slice of a third excitatory synapse. (l) Segmentation of the synapse in (k). (m) Zoom-in view of the synapse in (k). (n) IsoNet-corrected tomographic slice of (m). Red arrowheads indicate PSD nanoblocks. (o) Clustering results of 40-nm PSD density projected to the plane parallel to the postsynaptic membrane. Each color represents one cluster. Docked or tethered vesicle positions are marked as circles. (p) Distribution of PSD nanoblock area. (q) Distribution of nearest neighbor distance between PSD nanoblock centroids. (r) Distribution of distance between nearest pixels of PSD nanoblocks. (s-t) Distribution of normalized distance between potential release site and nanoblock. (u) Working model of subsynaptic organizations. Scale bars: (a, b, f, g, k, l) 100 nm; (c-e, h-j, m-o) 50 nm.

Furthermore, we observed multiple tethered vesicles, docked vesicles, or fused vesicles (Supplementary Fig. 4i-n) which serve as potential occupied release sites and are transsynaptically located in the boundary of PSD nanoblocks (Fig. 4s-t). In comparison to ∼1.8 potential release sites occupied by synaptic vesicles per synapse in synaptosomes, intact synapses in cultured neuron exhibit ∼4.3 potential release sites occupied by synaptic vesicles per synapse. Considering all docked, tethered, or fused synaptic vesicles (n=145) as potential release sites, we employed the same analysis of normalized vesicle-nanoblock centroid distances, and found no differences between the distribution of observed vesicle-nanoblock distances and those with randomized release sites (Fig. 1n). Like our previous results in synaptosomes, this further indicates that in hippocampal neuronal cultures the PSD nanoblocks are redundant for alignment with the potential release sites.

## Discussion

In this study, we utilized cryo-ET to directly visualize the ultrastructure of excitatory synapses in their near-native state of purified hippocampal synaptosomes from 10-week-old young adult rats as well as primary hippocampal neurons cultured from postnatal day 0 newborn rats. Importantly, we found that in our synaptosome samples, the exposed postsynaptic membranes remain attached to the presynaptic terminals. Thus, we were able to directly visualize membrane-associated PSD components in these preparations, which we refer as PSD nanoblocks. However, we acknowledge that some postsynaptic components located in the gap between PSD nanoblocks might be lost during sample preparation. This allowed us to study the PSD components against a relatively clean background. Meanwhile, studies of the whole synapse in intact cultured neurons provided more physiological and complete ultrastructure features. With these two different samples, we consistently captured a subsynaptic organization of PSD in the form of nanoblocks, which are facing presynaptic release sites. Our observations suggest that membrane-associated PSD components within synaptosomes are relatively stable, and that they maintain structural integrity during the isolation process for synaptosome preparation via our simplified protocol. This finding is consistent with previous observations that PSD complexes and the PSD membranes are detergent-resistant and stable enough to be purified from conventional synaptic fractions^39–42^. Thus, it is likely that the nanoscale organization and basic ultrastructure features we observed also reflect synaptic architecture *in vivo*.

An alignment between pre- and postsynaptic elements has been proposed but was only recently reported and named a *nanocolumn*, discovered through super-resolution light microscopy^3^. Thus far, only labeled synaptic proteins such as PSD95, receptors, and RIM have been shown to form nanoclusters as a part of nanocolumns. In contrast, our cryo-ET approach allows us to examine the molecular organization of all cellular components at molecular resolution, nanostructures not only those can be labeled for super-resolution light microscopy^31–33,51^. Martinez-Sanchez et al. applied cryo-ET to examine the alignment between presynaptic vesicles and postsynaptic receptors obtained through subtomogram averaging in synaptoneurosomes^10^. Here, we showed that the majority of the potential release sites, indicated by tethered, docked and fused vesicles, are transsynaptically located in the boundary of the PSD nanoblocks in both synaptosomes and intact neurons (Figs. 1 and 4). However, our analyses showed that this alignment is statistically indistinguishable from a random distribution of potential release sites and PSD nanoblocks. There might be several plausible explanations to this seemingly unreasonable result. First, the postsynaptic density might be redundant for neurotransmission, since majority of nanoblocks have less than 20 nm distance in between each other, encompassing the entire postsynaptic density in a near-continuous fashion. Second, technical limitations of sample preparation might have obscured a physiologically relevant alignment. For example, we have found fewer tethered, docked and fused vesicles in synaptosomes compared to intact neurons, suggesting that although the presence and distribution of postsynaptic nanoblocks are consistent throughout, the transsynaptic alignment might be susceptible to various steps in sample preparation. We also observed potential release sites that weren’t occupied by vesicles in synaptosomes (Fig. 1j-k), further confounding the analyses of transsynaptic alignment. Third and last, limitations of cryo-ET might have contributed to experimental absence of a specific transsynaptic alignment between release sites and PSD nanoblocks. For example, we were unable to further dissect the constituents of individual PSD nanoblocks, which might differ from one nanoblock to another, indicating distinct synaptic functions for each nanoblock requiring a specific transsynaptic alignment. Further improvements in cellular tomography will surely enable direct visualization of individual synaptic proteins *ex vivo* without requiring additional steps of sample preparation that might alter nano-scale ultrastructure of a synapse, providing with near native-state structures. Future advances in cryo-ET might reveal a specific alignment for different subgroups of postsynaptic nanoblocks with different release sites.

The PSD in the excitatory postsynaptic terminal is a dense area packed with postsynaptic proteins, where hundreds of molecules have been identified^52,53^. Efforts have been made to clarify the PSD’s molecular organization and the synaptic functions that these PSD proteins serve. The development of imaging techniques such as super-resolution fluorescent microscopy has facilitated the discovery of subsynaptic protein clusters, or *nanodomains*^3–5,8,54^. These nanodomains are ∼70 to 160 nm in size, which is larger than most protein nanoblocks we directly visualized within the PSDs of synaptosomes using cryo-ET. Therefore, we postulate that the small nanoblocks we discovered within the PSD may be a structural unit that forms the nanodomain, which in turn forms the PSD. A nanoblock consists of certain copies of several different scaffold proteins that form a supramolecular structure, which links to certain copies of synaptic receptors and allineates with the presynaptic release sites through adhesion molecules. Other postsynaptic proteins situated in the gaps between these nanoblocks may play a role in the regulation and organization of smaller nanoblocks into larger ones. However, considering the diversity of PSD protein complexes^55–58^, it remains possible that the composition of different-sized PSD nanoblocks varies. Further exploration of the composition of PSD nanoblocks will help us uncover the distinctions and similarities between nanoblocks and nanodomains.

Chemical fixation, or high-pressure freezing and freeze substitution EM sample preparation, displays a PSD mesh model^20,24,59^. Using cryo-ET, we also observed vertical filaments within synaptosomes and intact synapses, which were smaller in size. We suspect that the size difference is due to the inherent differences between sample preparation methods and imaging modalities. In the liquid condensate mode of PSD^28–30^, the liquid droplet is at micron scale. It is possible that the synaptic proteins in cells form nanoblocks or nanoclusters following the same mechanism as in the *in vitro* PSD protein mixing assay.

Subtomogram averaging of postsynaptic protein particles produced a 24-Å and a 26-Å structure from a small number of particles (Supplementary Fig.3). Our inability to achieve higher resolutions to identify protein composition is attributed to the complexity and diversity of postsynaptic proteins and protein complexes present on the postsynaptic membrane of brain tissue samples, which include various types of glutamate receptors^60–67^. The ligand proteins and neighboring proteins also contribute to the density, rendering each postsynaptic protein structurally heterogenous. Moreover, distinct postsynaptic receptors exhibit varying degrees of retention in biochemical PSD preparations^68–71^. Therefore, it is challenging to identify the protein complex based on the current resolution of our subtomogram averaging. When we compared the geometric shape of the average structure with existing PDB models, type A particles and type B particles seem to resemble the O-shaped and the Y-shaped AMPA receptors, respectively (Supplementary Fig. 3c-h). Similar to the comparison made by Martinez-Sanchez *et al.*^10^, our averaged structures don’t look like other available structures of postsynaptic complexes. It is likely the distribution of type A and type B particles shown in Fig. 3e-h reflected the distribution of different types of AMPA receptors in synapses. This is also consistent with studies showing AMPA receptor nano-organization does not fully align with PSD95 nano-organization^5,8,27^. Overcoming these difficulties in observing high-resolution structures of postsynaptic receptors, as well as linking their *in situ* conformation information with their functional states, would require an enormous cryo-ET dataset of intact synapses from cultured neurons or even brain tissues with proper thickness.

Here we propose a nanoscale organization model at excitatory synapses (Fig. 4u). The model is supported by our data from isolated synaptic terminals, and cultured neurons (Figs. 1-4), showing the commonality and diversity across synapses. Evidence from super-resolution and EM studies also suggests that synaptic activity affects the number and size of PSD nanoclusters or densities^4,72–74^. Thus, we postulate that the nanoblocks that we identified in both synaptosomes and intact neurons might be the fundamental unit of postsynaptic density and might play a role in regulating the synaptic strength.

## Acknowledgments

We thank Drs. Ege T. Kavalali, Terunaga Nakagawa, William Wan, Roger J Colbran and Mark E. Bowen for stimulating discussions and critical reading of the manuscript; the NIH for support (R00 MH113764 to Q.Z., R01 MH132918-01A1 to Q.Z.); the Vanderbilt Faculty Fellowship Endowment Fund; and “Neurodegenerative” TIPS Initiative Award for support.

EM data collections were conducted at the Center for Structural Biology Cryo-EM Facility at Vanderbilt University. We also acknowledge the use of the Glacios cryo-TEM, which was acquired by NIH grant 1 S10 OD030292-01.

## Author contributions

Conceptualization, R.S. and Q.Z.; Methodology, R.S. and Q.Z.; Investigation, R.S. and Q.Z.; Experiments, R.S. and B.A., Data analysis, R.S., J.P.A., L.W., M.H., X.W. and ZM.Z.; Writing – Original Draft, R.S. and Q.Z.; Writing – Review & Editing, R.S., J.P.A., L.W., M.H., X.W., ZM.Z. and Q.Z..; Funding Acquisition, Q.Z..

## Methods

### Synaptosome isolation

10-week Sprague-Dawley rats with either sex were used for synaptosome isolation. The rats were kept in 12 hours: 12 hours dark:light cycle and provided with treats as well as cardboard enrichments before the experiments. The rats were deeply anesthetized with isoflurane. Hippocampi tissue were dissected in the ice-cold homogenization buffer containing 0.32 M sucrose, 4 mM HEPES, 1mM EDTA, at pH 7.4 with or without protease inhibitor cocktail (1 tablet in 50 ml homogenization buffer) and 20 mM DTT. Then the tissue was homogenized at ∼500 rpm for 4 strokes using an overhead stirrer (Electron Microscopy Sciences 6480610) and a tissue grinder (Electron Microscopy Sciences 6479310) in the ice-cold homogenization buffer. After homogenization, the homogenates were centrifuged at 800g at 4 ℃ for 10 min. The supernatant was immediately taken for cryo-ET sample preparation. All animal procedures were performed in accordance with the guide for the care and use of laboratory animals and were approved by the Institutional Animal Care and Use Committee at Vanderbilt University.

### Primary dissociated neuronal cultures

The protocol was modified from previous works^22,75,76^. Postnatal day 0 Sprague-Dawley rats of either sex were used for primary hippocampal cultures. Pregnant Sprague-Dawley rats were housed individually until they gave birth to a litter and were kept in 12 hours: 12 hours dark:light cycle. The pregnant rats were provided with the same treats as well as cardboard enrichments. Postnatal day 0 littermates were used to prepare primary dissociated neuronal cultures. Hippocampi were dissected in ice cold 20% fetal bovine serum (FBS) containing Hanks’ balanced salt solution. Tissues were then washed and treated with 10 mg/ml trypsin and 0.5 mg/ml DNase at 37 ℃ for 10 min. The tissues were washed again, dissociated using filtered P1000 tip and centrifuged at 1000 rpm for 10 mins at 4 ℃. Pellet containing neurons was resuspended in Neurobasal Plus medium supplemented with GlutaMAX-I and B27 supplement. Neurons were plated onto Quantifoil R2/2 Au 200 EM grids coated with poly-L-lysine. Cultures were kept in humidified incubators at 37 ℃ and gassed with 95% air and 5% CO2. On DIV1, the neurobasal plus medium was replaced with 4 μM cytosine arabinoside containing neurobasal plus medium. On DIV4, the cytosine arabinoside concentration was dropped 2 μM by performing a half media change. Cultures were kept without any disruption until DIV14. The cultures on EM grids were plunged frozen between DIV14-18, when synapses reached maturity. All animal procedures were performed in accordance with the guide for the care and use of laboratory animals and were approved by the Institutional Animal Care and Use Committee at Vanderbilt University.

### Synaptosome and neuronal culture vitrification

#### Synaptosome

FEI Vitrobot Mark III was used for plunge freezing. Set the parameters of the plunge freezer as follows: humidity 100%, temperature 4 ℃, blot time 3.5 s, blot total 1, blot offset -1.5, and wait time 6 s. Supernatant after centrifugation mixed with 10-nm gold beads solution (Aurion, cat # 25486) at 1:1 ratio was added to the mounted grid (Quantifoil R2/2 Cu 200 EM). No other chemicals or stimulation were applied prior to plunge freezing. The grids were plunged into liquid nitrogen cooled liquid ethane for rapid vitrification and were stored in liquid nitrogen until use.

#### Primary dissociated neuronal cultures

The vitrification process of neuronal cultures is similar to the synaptosome with a few modifications. The culture dishes were taken out from the incubator on DIV 14-18. The culture medium was replaced with a modified Tyrode’s solution containing the followings was used: (in mM): 150 NaCl, 4 KCl, 1.25 MgCl_2_, 2 CaCl_2_, 10 D-glucose, 10 HEPES at pH 7.4. Then the cultured neurons were kept in the modified Tyrode’s solution for 10min. After setting up the plunge freezer as follows: humidity 100%, temperature 22 °C, blot time 3.5 s, blot total 1, blot offset -1.5, and wait time 6 s, the grids with cultures were mounted. 4 μL 10-nm gold beads solution (in the modified Tyrode’s solution) was added to the mounted grid before plunging. No other chemicals or stimulation were applied prior to plunge freezing. After rapid vitrification, the grids were stored in liquid nitrogen until use.

### Cryo-ET data collection

Automated batch tilt series collection was done with Thermo Fisher Tomography5 software on Thermo Fisher Titan Krios G4 using the Gatan K3 camera in zero-loss mode (slit width 20 eV). The tilt series of neuronal cultures were collected using the dose-symmetric scheme^77^ starting from 0° to ± 60° with an interval of 2°, and with the defocus value at -7 μm, with the total electron dosage of ∼150 e^−^/Å^2^ which were evenly distributed between tilts. The final pixel size was 3.3 Å with 26000× magnification. The tilt series of synaptosomes were collected with Voltage Phase Plate. The phase plate dataset was collected at the pixel size 3.3 Å with 26000× magnification, using the dose-symmetric scheme starting from 0° to ± 60°with an interval of 2°, and with the defocus value at -2 μm. The total electron dosage of ∼150 e^−^/Å^2^ was evenly distributed between tilts. The phase plate was switched to a new position and activated to gain a phase shift of around 0.3π prior to each tilt series. Conditioning of the phase plate was also performed between each tilt.

### Data reconstruction and subtomogram averaging

Tomograms of synaptosome at pixel size of 3.3 Å with phase plate were used for subtomogram averaging. Synaptosome thicker than 500 nm were excluded, and 28 tomograms were selected. TOMOMAN package^78^ was used to preprocess the data. The workflow of TOMOMAN included motion correction by MotionCorr2^79^, image sorting, dose filtering and CTF estimation by Gctf ^80^. Then the tilt series were aligned the reconstructed using IMOD^81^. Fine alignment of the tilt series was performed by using the 10 nm gold beads (Aurion, cat # 25486) as fiducial markers. Both back projection and back projection with five-iteration SIRT (simultaneous iterative reconstruction technique)-like were performed to reconstruct each tilt series. Segmentation and 3D rendering were done using IMOD. In the synaptosome dataset, 1565 particles were picked according to the size and shape. Only the particles around 10 ∼14 nm on the postsynaptic membrane were included. Subtomogram averaging was done using RELION 3.0.8^82^. We applied a box size of 96 using unbinned data. First, we generated the initial model using 3d auto-refine with the unbinned data without applying symmetry. 3d classification resulted in two good classes. After 3d classification, we generated 3d initial models for each class respectively with C2 symmetry and did 3d auto-refine for each class using the corresponding initial models. Manual repicking of the particles was performed to exclude the particles which were too close (distance < 8 nm) and final averaging was performed using the remaining particles in 3d auto-refine.

### PSD density analysis and quantification

Density of PSD was measured using the following method. In the synaptosome dataset, synaptosome with a PSD close to cryo grid carbon hole were excluded as the gray values were affected by the hole edge. Synaptosomes thicker than 500 nm and synaptosomes with the postsynaptic membrane not parallel to its presynaptic membrane were also excluded. 123 tomograms (58 tomograms for synaptosome isolated without protease inhibitor and DTT in the homogenization buffer, 41 tomograms for synaptosome isolated with protease inhibitor and DTT in the homogenization buffer, and 24 tomograms for synaptoneurosomes isolated with and without protease inhibitor and DTT in the homogenization buffer) were selected for PSD density clustering analysis. In the culture neuron dataset, synapses thicker than 700 nm were excluded, and 34 tomograms were selected for PSD density analysis. The gray values within 40 nm range from the postsynaptic membrane were projected to xz plane (Fig. 2) using Fiji^83^. The gray value of each pixel of the projected 2D plane was the average value of the 40 nm column. Next, we used different strategies to process synaptosome and cultured neuron dataset. For synaptosome data, we measured the average gray value in empty space as background gray value. Then we chose the bottom 25% gray values (25% darkest) below the background value for the following clustering analysis. For synapses from cultured neurons, we analysed the density in the IsoNet^50^ corrected tomograms. The data set was normalized after IsoNet internal process, and we chose the bottom 20% gray values (20% darkest) for the following clustering analysis. Lastly, clustering analysis algorithm DBSCAN (density-based spatial clustering of applications with noise) was adopted to do the clustering of PSD density. In the synaptosome sample analysis, DBSCAN with different parameters were performed (Supplementary Fig. 2) and DBSCAN (6, 32) was applied as the results are consistent with the visualized electron density clusters in the tomographic slices. In the synaptosome isolated with protease inhibitor and DTT, synaptoneurosome and synapse analysis, DBSCAN (7, 32) was applied.

The area of a nanoblock was defined as its convex hull area shown by red lines in Fig. 2b-c and Supplementary Fig. 2b bottom panel. All area distributions were cut off to 3500 nm^2^ or 6000 nm^2^ to show the fitting peaks more clearly. The width of a nanoblock was defined as the largest x-axis distance between two pixels in the cluster. The widths of nanoblocks are also marked in Figures 2c and the bottom panel of 2d. To precisely measure the height of a nanoblock, we first averaged the tomographic slices in the range that contains the nanoblock determined in the DBSCAN clustering results. Then we drew a line as shown in Fig. 1c to measure the gray value of the nanoblock in y axis at its largest length. The height of the nanoblock was defined as the distance from the farthest pixel of the nanoblock to the postsynaptic membrane. Since we took the weight of each pixel in the nanoblock the same, the centroid of a nanoblock is the geometry centroid of the nanoblock by averaging all coordinates of the pixels of a nanoblock. The distance between the nanoblock edge and centroid of the nanoblock was defined as the distance from the farthest pixel of the nanoblock to centroid of the nanoblock.

Positions of all tethered, docked, or fused vesicles were projected to the xz plane in the same coordinate system of PSD density to analyse the relation between potential release sites and PSD nanoblocks for both synaptosomes (vesicle number is 106) and synapses (vesicle number is 145). d/D was calculated as shown in Fig. 1l and 4t. Here d was defined as the distance from synaptic vesicle centroid to nanoblock centroid. D was defined as the distance from farthest pixel of the nanoblock to centroid of the nanoblock. Thus d/D is the normalized distance to from the synaptic vesicle to the nanoblock. To test if the alignment between potential release sites and PSD nanoblock is specialized, the position of the vesicles were randomized within the bounds of the entire PSD. We randomized each synaptic vesicle position one hundred times, and then performed the analysis for the randomized vesicle position to calculate d/D as we did for actual vesicles. Positions of postsynaptic membrane proteins were also projected to the xz plane in the same coordinate system of PSD density to analyse the relation between membrane proteins and PSD nanoblocks. Then we calculate d/D for each protein particle. Here d was defined as the distance from particle to nanoblock centroid. D was defined as the distance from farthest pixel of the nanoblock to centroid of the nanoblock.

### Data and code availability

All data and code reported in this paper will be shared by the lead contact upon request.

---

**Supplementary Fig. 1.**
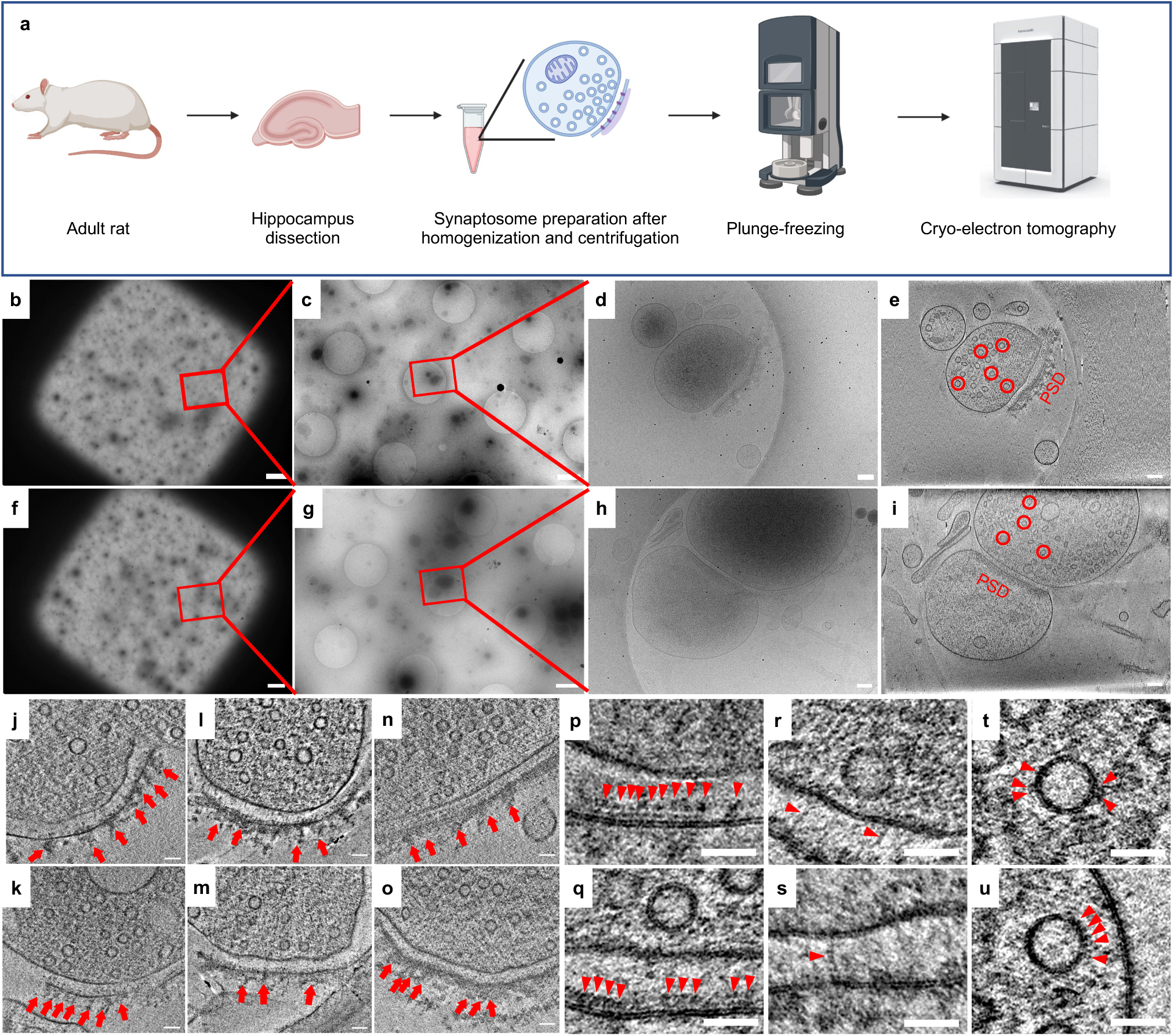

**Supplementary Fig. 2.**
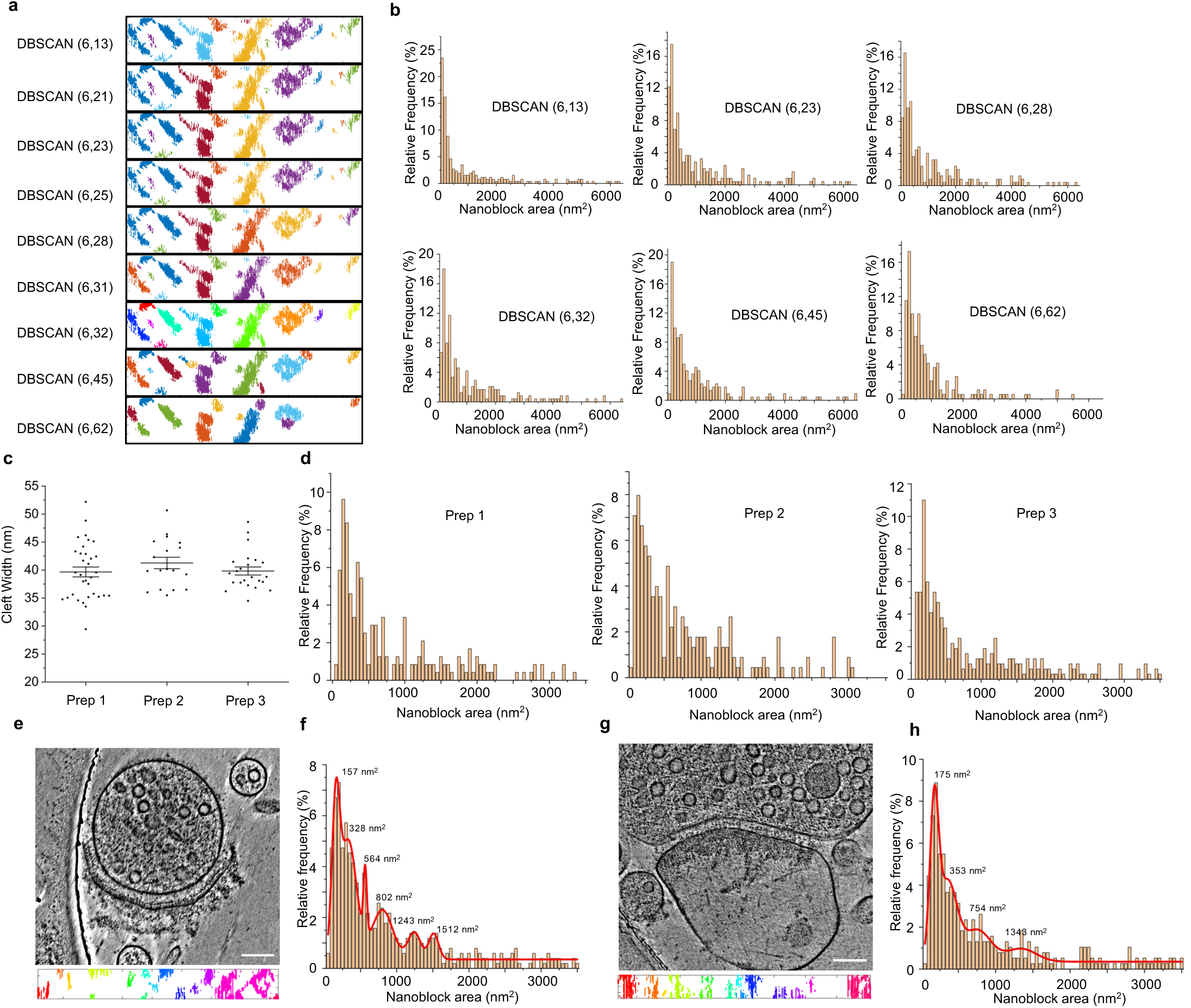

**Supplementary Fig. 3.**
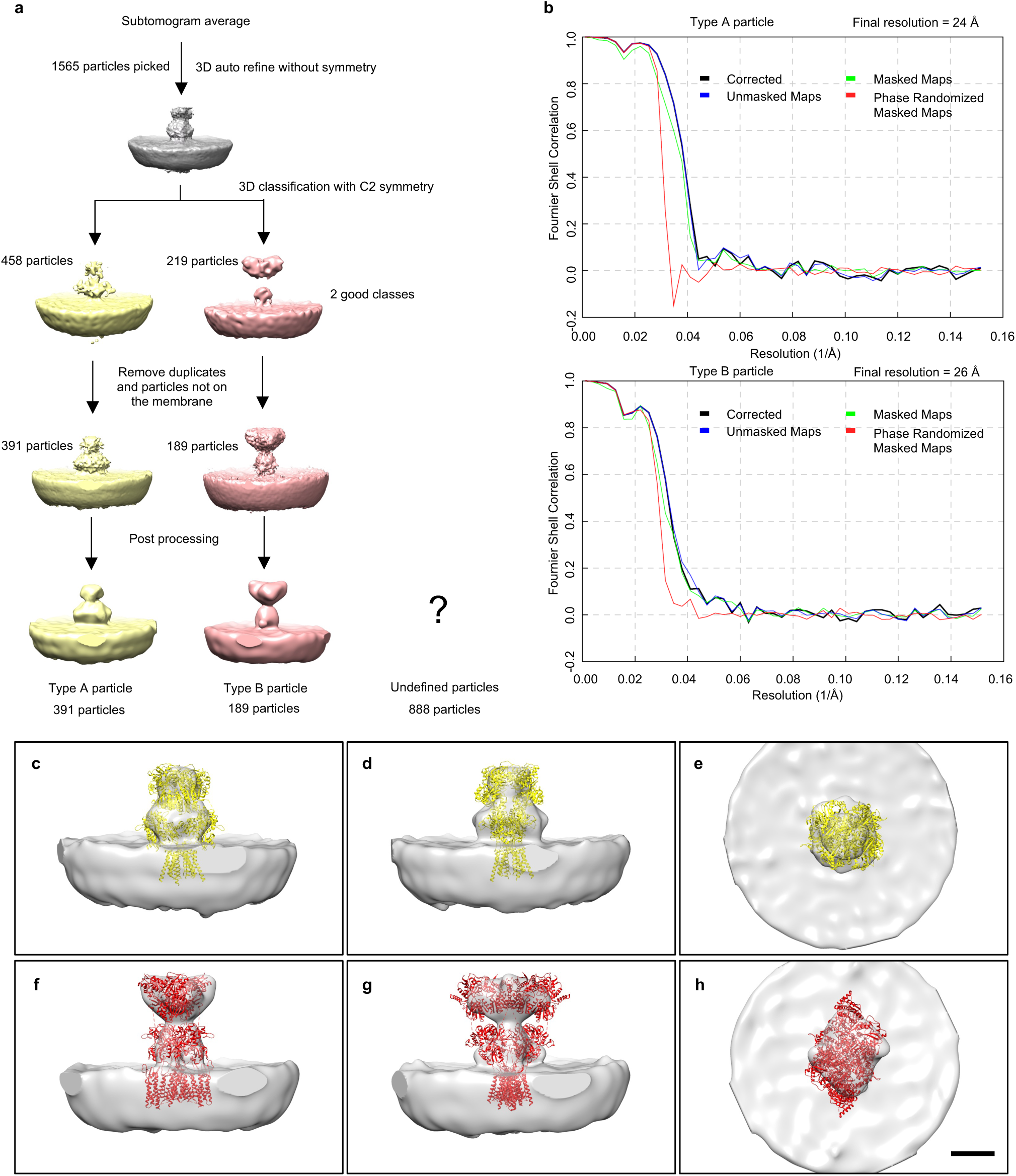

**Supplementary Fig. 4.**
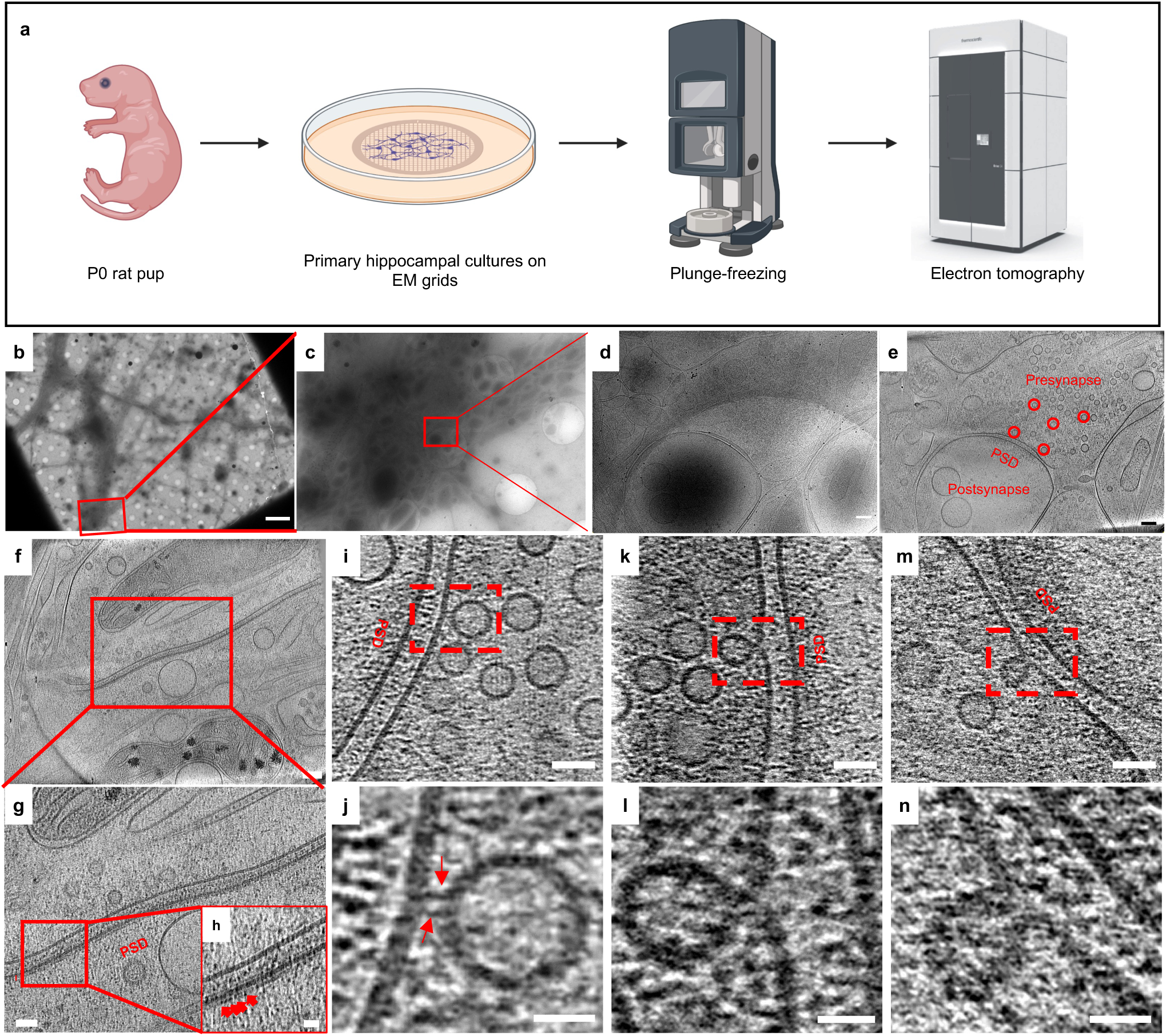

