## Supplementary_documents for "Cryo-electron tomography reveals postsynaptic nanoblocks in excitatory synapses": Supplementary_information_v13_clean.pdf

**Supplementary Fig. 1**. Cryo-electron tomography of synaptosome with postsynaptic membrane and PSD. (a) Workflow of imaging synaptosomes with PSD on gold EM grids using cryo-electron tomography. (b-d) Images of a synaptosome with PSD at different magnifications in cryo-EM. (e) shows a virtual slice of the reconstructed tomogram of the same area in (d). A synaptosome containing synaptic vesicles indicated by red circles and postsynaptic density (PSD). (f-h) Images of a synaptosome with enclosed postsynapse at different magnifications in cryo-EM. (i) shows a virtual slice of the reconstructed tomogram of the same area in (h). A synaptosome containing synaptic vesicles indicated by red circles and postsynaptic density (PSD) in an enclosed postsynapse. (j, k) Synaptosomes with separate PSD nanoblocks. (l, m) Synaptosomes with separate PSD nanoblocks and continuous PSD. (n, o) Synaptosomes with continuous PSD. Arrows indicate PSD nanoblocks. Tomographic slices show membrane particles (p, q), adhesion molecule-like particles (r, s) and synaptic vesicle associated proteins (t, u) as arrowheads indicate in synaptosomes. Scale bars: (b, f) 5 µm; (c, g) 1 µm; (d, h) 100 nm; (e, i) 100 nm; (j-u) 50 nm.

**Supplementary Fig. 2**. DBSCAN analysis. (a) DBSCAN clustering results of an excitatory synaptosome PSD with different DBSCAN parameters. (b) Distribution of nanoblock area with different DBSCAN parameters. (c) Cleft widths of synaptosomes in three preparations isolated without protease inhibitor and DTT. (d) Distribution of nanoblock area in three preparations isolated without protease inhibitor and DTT. (e) A tomographic slice showing an excitatory synaptosome isolated with protease inhibitor and DTT, and corresponding DBSCAN clustering results of PSD density. (f) Distribution of nanoblock area of synaptosome isolated with protease inhibitor and DTT. (g) A tomographic slice showing an excitatory synaptoneurosome and corresponding DBSCAN clustering results of PSD density. (h) Distribution of nanoblock area of synaptoneurosome combined from all isolation conditions.

**Supplementary Fig. 3**. Subtomogram averaging of membrane proteins from synaptosomes. (a). Steps for subtomogram averaging. (b). Gold-standard FSC shows the final resolution of sub-tomogram averaging structures of type A particle and type B particle. (c-h) Subtomogram averaged type A particle (c-e) and type B particle (f-h) with AMPA receptor model fitting. Models from PDB bank are fitted to the averaged electron densities. Yellow: PDB 5ide. Red: PDB 6qkz. Scale bar: 5nm.

**Supplementary Fig. 4**. Cryo-electron tomography of primary cultured neurons. (a) Workflow of imaging primary cultured neurons on gold EM grids using cryo-electron tomography. (b-d) Images of neurons at different magnifications in cryo-EM. (e) shows a virtual slice of the reconstructed tomogram of the same area in (d). There is a synapse containing synaptic vesicles indicated by red circles and thick postsynaptic density (PSD). (f-g) The tomogram slice shows a synapse with thin PSD. Arrows in (h) indicate the proteins in the PSD. (i-n) Synaptic vesicles at different stages of exocytosis show release sites in the presynaptic membrane. (i, j) Tethered vesicle that tethers to the presynaptic membrane with linkers. (k, l) Docked vesicle with membrane directly contacting presynaptic membrane. (m, n) Fused vesicle with vesicular membrane fused to presynaptic membrane. Scale bars: (b) 5 µm; (c) 1 µm; (d) 100 nm; (e) 100 nm; (f)100 nm; (g) 50 nm; (h) 20 nm; (i, k, m) 50 nm; (j, l, n) 20 nm.

**Supplementary Video 1.** 3D surface rendering of a synaptosome observed by cryo-electron tomography with putative type A and type B particles anchored on the postsynaptic membrane. Spherical structures (cyan) are synaptic vesicles. Other subcellular structures are presynaptic membrane (green), postsynaptic membrane (deep blue), postsynaptic density (gold), type A particles (yellow) and type B particles (red). Scale bar: 100 nm.

**Supplementary Video 2.** 3D surface rendering of a synapse in primary cultured neurons observed by cryo-electron tomography. Spherical structures (cyan) are synaptic vesicles. Other subcellular structures are presynaptic membrane (green), postsynaptic membrane (deep blue), postsynaptic density (gold), microtubules (light blue), endoplasmic reticulum (purple), presynaptic endosomes (red) and postsynaptic endosome (red). Scale bar: 200 nm.
